# Reversibility of chemotherapy-induced senescence is independent of autophagy and a potential model for tumor dormancy and cancer recurrence

**DOI:** 10.1101/099812

**Authors:** Tareq Saleh, Emmanuel K. Cudjoe, S. Lauren Kyte, Scott C. Henderson, Lynne W. Elmore, David A. Gewirtz

## Abstract

Autophagy and senescence are both well-established responses to chemotherapy and radiation that often occur in parallel, contributing to growth arrest in tumor cells. However, it has not been established whether this growth arrest is reversible. This question was addressed using non-small cell lung cancer models exposed to the cancer chemotherapeutic drug, etoposide. Senescent cells that were sorted, identified by β-galactosidase staining and alterations in morphology, isolated by flow cytometric cell sorting based on C_12_FDG staining, and real-time live microscopy were found to be capable of recovering proliferative capacity. Autophagy, monitored by vacuole formation, SQSTM1/p62 degradation, and LC3BII generation did not interfere with either the senescence arrest or proliferative recovery and was nonprotective in function (i.e. autophagy inhibition via both pharmacological and genetic strategies had negligible impact on the response to etoposide).

These observations argue against the premise that (chemotherapy-induced) senescence is irreversible and indicate that therapy-induced senescence may ultimately be a transient process in that at least a subpopulation of tumor cells can and will remain metabolically active and recover proliferative capacity independently of autophagic turnover. We therefore propose that dormant tumor cells may be capable of prolonged survival in a state of autophagy/senescence and that disease recurrence may reflect escape from this senescence-arrested state.

## Introduction

Autophagy and senescence are both well-established responses to cellular stress^1,2^. Autophagy and senescence, whether induced by oncogene activation, chemotherapy or radiation often, if not always, occur in parallel, although we and others have determined that these responses are dissociable (i.e. senescence occurs and/or is sustained even when autophagy is suppressed)^3–5^.

Autophagy, which is predominantly cell survival mechanism, has been proposed to contribute to therapy resistance and potentially to tumor dormancy^1,6^. With regard to tumor dormancy, this would require that some subset of the dormant tumor cell population could eventually escape from the dormant state. In contrast, senescence, whether a consequence of telomere shortening (replicative senescence), oncogene activation (oncogene-induced) or that which occurs in tumor cells in response to chemotherapy or radiation (therapy-induced senescence, accelerated or premature senescence) is generally considered to be irreversible ^7–10^. However, it is clear that senescent cells actually retain reproductive potential since transformation and immortalization involve escape from replicative senescence ^11–13^. A recent review published by our research group highlights studies of the capacity of cells to escape from the various forms of senescence^14^. With regard to therapy-induced senescence, reports by our group^15,16^ as well as other laboratories have presented evidence in support of the premise that tumor cells in a state of senescence are not obligatorily in a terminally growth-arrested state ^17–26^. These studies generally involved genetic manipulation of cell cycle regulators, such as p53 or p16^INK4a^; however, it has remained uncertain whether spontaneous restoration of proliferative capacity can occur from the senescent state. Furthermore, conclusions relating to the reversibility of senescence have generally been based on studies in mass culture, where the origin of the replicating cells cannot be unequivocally attributed to the senescent population.

A primary finding of the current work is the capacity of etoposide-induced highly autophagic, senescent lung cancer cells, to resume cellular division, which we have previously termed *proliferative recovery* ^2^. Furthermore, autophagy and senescence induced by etoposide are found to be dissociable and the capacity of the cells to recover post-drug treatment is not compromised when autophagy is inhibited either pharmacologically or genetically. Finally, the autophagy is clearly *non-cytoprotective* in function, as we have previously shown in other experimental models ^15,27,28^.

Based on these findings, we propose the novel premise that therapy-induced senescence is reversible and independent of autophagic cell survival. Senescence, rather than autophagy, is likely to play a critical role in determining the nature of the tumor cell response in terms of transient growth arrest, the relative lack of cell death, and the ultimate re-emergence of the tumor cells from the growth-arrested state to regain proliferative function and contribute to disease recurrence.

## Results

### Etoposide promotes growth arrest, autophagy and senescence in NSCLC cells followed by proliferative recovery

H460 human NSCLC cells undergo growth arrest followed by proliferative recovery upon exposure to etoposide, a mainstay chemotherapeutic agent (**Figure 1A, left panel**). Exposure to 1 μM etoposide, the concentration utilized in all subsequent studies, resulted in growth arrest for at least five days followed by proliferative recovery at approximately 7 days post-drug exposure. A similar response pattern to etoposide was also observed in the A549 NSCLC cell line (**Figure 1A, right panel**). Recovery of H460 cells is also shown in the colony forming assay in **Figure 1B**. As would have been anticipated based on the fact that etoposide promoted autophagy in A549 and U1810 NSCLC cell lines^29^, autophagy was also evident in the H460 cells exposed to etoposide. **Figure 1C, left panel,** shows the concentration-dependent formation of acridine orange-stained acidic vesicular organelles in the H460 cells 48 hours following removal of drug (also quantified by flow cytometry **Figure 1C**). The induction of autophagy was confirmed based on the increased formation of GFP-LC3 puncta in H460 cells (**Figure 1D**), as well as the increased lipidation of LC3 (conversion of LC3 from form I to form II) and the degradation of p62/SQTSM1 (**Figure 1E**), indicating that the autophagic process is progressing to completion (i.e. autophagic flux).

**Figure 1.**
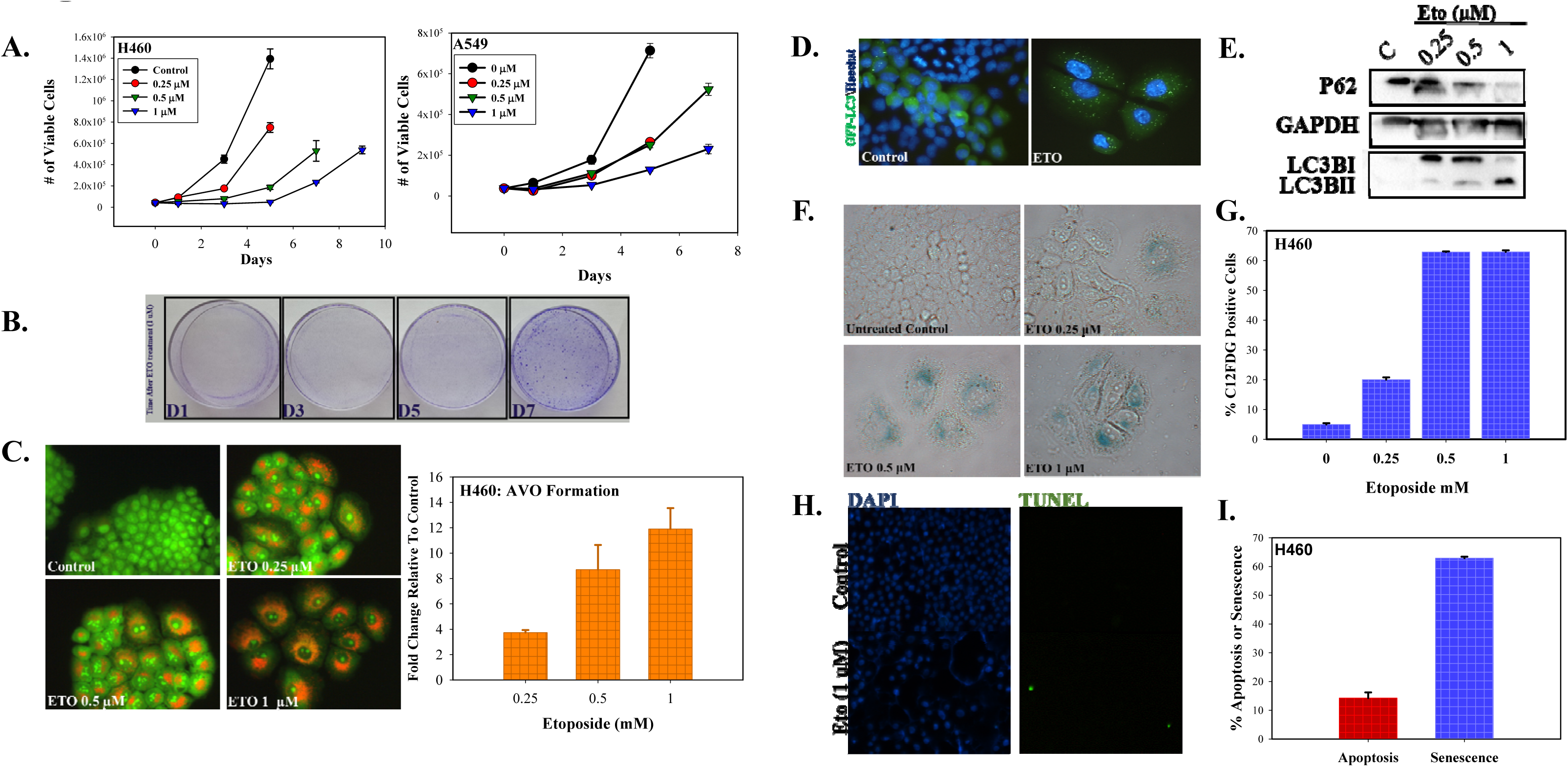
Etoposide promotes growth arrest, autophagy and senescence in NSCLC cells followed by proliferative recovery. **A.** H460 and A549 cells were exposed to etoposide at concentrations of 0.25, 0.5, or 1 μM for 24 h (day 0); cell number was determined over the indicated time points. (n=3). **B.** Colony formation of H460 cells following 1μM etoposide exposure for 24 h. (n=2) **C.** Fluorescence microscopy showing concentration-dependent increase in acridine orange-stained vacuoles induced by 0.25, 0.5, and 1 μM etoposide and quantification of acridine orange staining by FACS analysis in response to increasing concentrations of etoposide (n=2). Imaging and FACS were performed 48 h after drug removal. (20x objective). **D.** Fluorescence microscopy showing increased GFP-LC3 puncta in response to 1 μM etoposide exposure. Imaging performed 48 h after drug removal. (20x objective) (n=2) **E.** Western blot showing the dose-dependent decrease in p62/SQSTM1 and the lipidation of LC3B in response to etoposide (n=2). **F.** Senescence-associated β-galactosidase staining of H460 cells exposed to 0.25, 0.5, or 1 μM etoposide (20x objective). **G.** Quantification of senescence based on C_12_FDG staining of H460 cells followed by FACS analysis. Staining and analysis were performed 48 h after drug removal (day 3) (n=3). **H.** TUNEL assay of H460 cells 48 h following etoposide (1μM) removal. **I.** Percentage of apoptotic H460 cells following # h exposure to 1 μM etoposide was determined by Annexin V immunolabeling and FACS at the same time point. Data are expressed as standard error of the mean (SEM) for “n” independent experiments.

Furthermore, and as early as 3 days after the initiation of drug exposure, H460 cells exhibit numerous features collectively indicative of senescence, specifically a flattened and enlarged appearance with abundant granulation, and histochemical staining for SA-β-galactosidase (SA-β-gal) activity (**Figure 1F**). Flow cytometric analysis quantifying the percentage of H460 cells expressing SA-β-gal activity at different concentrations of etoposide, relying on an established C_12_FDG fluorescent labeling procedure^30^, is presented in **Figure 1G**. In contrast to the promotion of senescence and autophagy, 1 μM etoposide produced minimal apoptosis in the H460 cells as shown by TUNEL assay (**Figure 1H**) and quantification of Annexin V-positive cells by flow cytometry (**Figure 1I**). These data collectively suggest that etoposide induces a senescent growth arrest in H460 cells associated with robust autophagic vacuolation. Footnote 1.

### Autophagy plays a non-cytoprotective role in response to etoposide in H460 cells and does not interfere with the ability of cells to recover proliferative capacity

Autophagy has historically been considered a survival response under conditions of nutrient deprivation or hypoxia, as well as a process that facilitates tumor growth and serves as a mechanism of resistance to therapy^31,32^. To determine whether autophagy might be protecting the H460 cells against etoposide toxicity and/or facilitating recovery following the senescent growth arrest, autophagy was suppressed using both pharmacological and genetic strategies, and the impact on sensitivity to etoposide was monitored. Cells were pretreated for 3 hours with the autophagy inhibitors chloroquine (CQ, 10 μM) or bafilomycin A1 (Baf, 5 nM) followed by 24 hours of exposure to etoposide in the presence of the CQ or Baf. The presence of CQ and Baf resulted in failure of lysosomal acidification, which is reflected by the yellow staining of vacuoles by acridine orange (**Figure 2A**); autophagy inhibition was confirmed by decreased degradation of p62/SQTSM1 in the presence of CQ or Baf (**Figure 2B**). **Figure 2C** shows that inhibition of autophagy did not alter sensitivity to etoposide in both MTT and clonogenic survival assays (except moderately with Baf at 1 μM etoposide), indicating that the autophagy was not cytoprotective. This conclusion relating to the function of autophagy was supported by the temporal response studies presented in **Figure 2D** where neither CQ nor Baf was able to alter the profile of growth arrest; furthermore, proliferative recovery occurred in the cells treated with etoposide both in the absence and presence of CQ or Baf.

**Figure 2.**
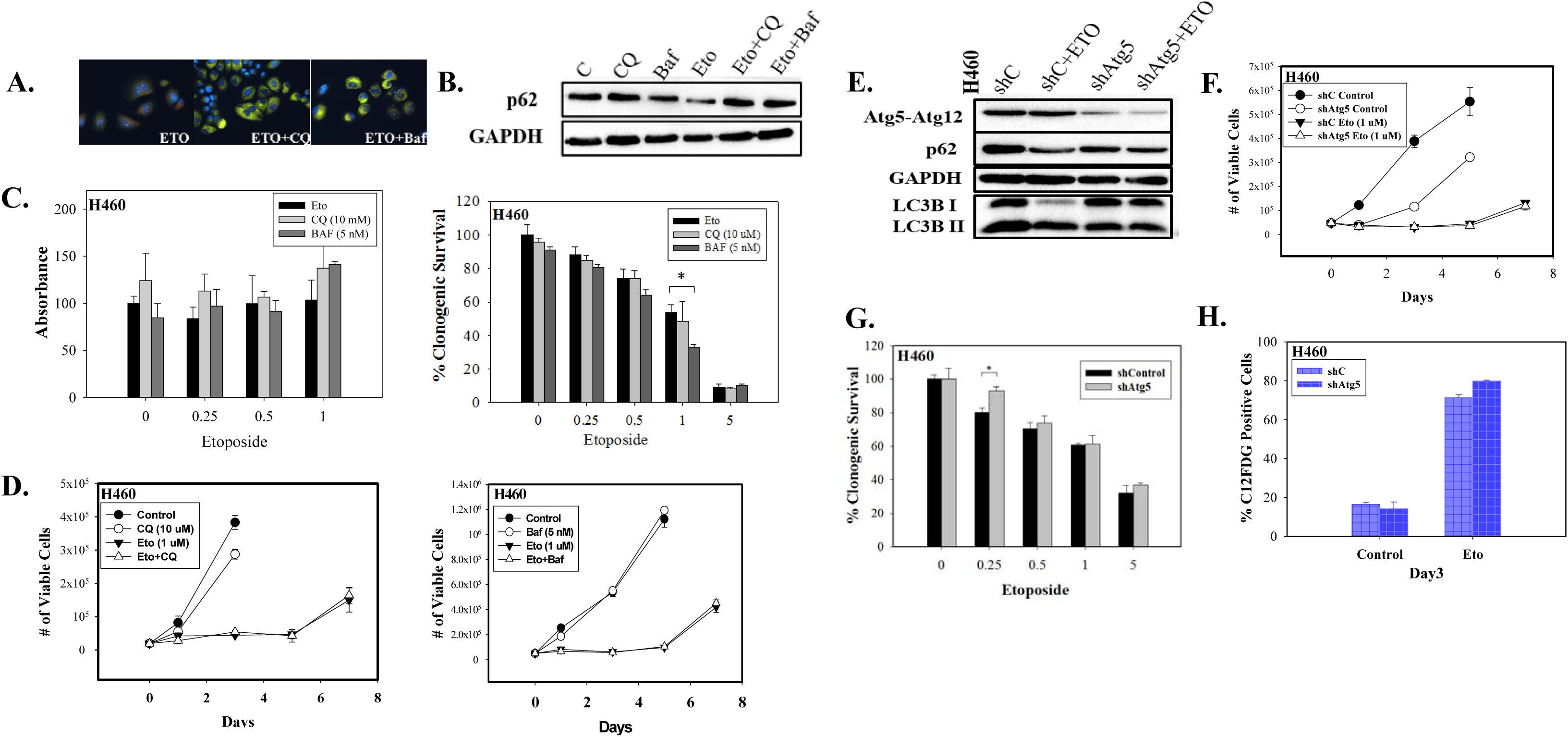
Autophagy plays a “non-cytoprotective” role in response to etoposide in H460 cells and does not interfere with the ability of cells to recover proliferative capacity. **A.**Fluorescence microscopy showing failure of lysosomal acidification following CQ (10 μM) or Baf (5 nM) co-treatment with 1μM etoposide. Cells were pretreated with CQ and Baf followed by an additional 24 h with etoposide. Images were taken 48 h after drug removal (20x objective). Nuclei stained with Hoechst 33342. (n=2). **B.** Western blot showing autophagy blockade by CQ (10 μM) and Baf (5 nM) based on levels of p62/SQSTM1 (n=3). C. ***Left panel.*** MTT assay showing influence of CQ (10 μM) or Baf (5 nM) on sensitivity of H460 cells to etoposide. Cells were pretreated with CQ or Baf for 3 h followed by co-treatment with etoposide for 24 h. Absorbance was measured 72 h after drug removal and replacement with fresh medium (n=3). Bars represent mean ± SD absorbance values relative to untreated control. C. **Right panel.** Clonogenic survival assay showing influence of CQ (10 μM) or Baf (5 nM) on sensitivity of H460 cells to etoposide (n=2). Cells were pretreated with CQ or Baf for 3 h followed by co-treatment with etoposide for 24 h. Colonies were counted 7 days following removal of drugs and replacement with fresh medium. Bars represent mean survival ± SD relative to untreated controls (α=0.05/3, *\* p<0.016).* **D.** Viable H460 cell number was determined at the indicated days following etoposide exposure in combination with CQ (10 μM) or Baf (5 nM) (n=2). **E.** Western blot following shRNA-mediated knockdown of *Atg5* (n=2). **F.** Temporal response to etoposide in parental H460 cells and H460 cells with knockdown of *Atg5* (n=2). **G.** Clonogenic survival assay comparing sensitivity of shControl and shAtg5 H460 cells in response to multiple etoposide concentrations. Bars represent mean survival ± SD relative to untreated controls (**α**=0.05/3, * *p<0.016)* (n=2). **H.** Percent senescence based on C_12_FDG staining at day 3 post-etoposide exposure in shControl cells and cells infected with shAtg5. Unless stated otherwise, data are expressed as standard error of the mean (SEM) for “n” independent experiments.

Autophagy was also blocked genetically using shRNA-meditated knockdown of *Atg5.* Confirmation of gene silencing is shown by immunoblotting where p62/SQTSM1 levels are increased and LC3BI to LC3BII conversion is reduced (**Figure 2E**). **Figure 2F** shows that growth curves were similar in H460 cells where *Atg5* was silenced as in the autophagy-competent cells. Furthermore, silencing of *Atg5* did not sensitize the H460 NSCLC cells to etoposide (except for a small effect at the 0.25 μM concentration, **Figure 2G**). In addition, inhibition of autophagy using both pharmacologic and genetic approaches did not significantly alter the extent of cell death induced by etoposide, as determined by Annexin/PI staining and FACS analysis (data not shown). It is therefore concluded that etoposide-induced autophagy in H460 cells is non-cytoprotective in function and its inhibition is not associated with increased cell death, consistent with the data presented in **Figures 2D and F**. These observations lead to the conclusion that autophagy plays a minor, if any, role in facilitating proliferative recovery in this system or interfering with the fate of the senescent cells.

In previous studies of radiation-induced autophagy and senescence in HCT-116 colon carcinoma cells, we demonstrated a direct correspondence between the extent of autophagy and senescence, where both responses were related to the extent of DNA damage ^15^. However, in this work as well as studies of doxorubicin-induced autophagy and senescence in breast tumor cells, we report that the two responses are clearly dissociable^5,15^. Our studies further demonstrate that the senescence induced by etoposide in the H460 cells is *independent* of autophagy, as genetic silencing of autophagy failed to influence the promotion of senescence by etoposide (**Figure 2H**).

### Proliferative recovery is associated with a decline in SA-β-galactosidase activity, entry into G1 and reduction in p21^Waf1^ expression

A time course analysis following exposure of H460 (and A549 cells) to 1 μM etoposide for 24 hours, based on the percentage of C_12_FDG-positive cells (Figure 3A), coincides with the profile of growth arrest and proliferative recovery observed in **Figure 1**. Cell cycle analysis confirmed the transient accumulation of H460 cells in the G2 phase of the cell cycle followed by their re-entry into the G1 phase at approximately the same time where we consistently observe the restoration of growth capacity (**Figure 3B**). It is furthermore of importance to note that a fraction of the cell population was polyploid, which has been suggested to be required for the generation of daughter cells from senescent precursors^17,18,33^. A largely similar pattern is evident for both mRNA and protein levels of p21^Waf1^ (**Figures 3C and D**), as well as the expression of IL-6, a component of the senescence-associated secretory phenotype (SASP) that is strongly associated with DNA-damage-induced senescence^34^ (**Figure 3D**). Interestingly, while DNA damage is elevated at day 1 post-drug exposure and declines over time (based on phosphorylated γH2AX levels), DNA integrity is not fully restored even by day 7 (**Figure 3E**).

**Figure 3.**
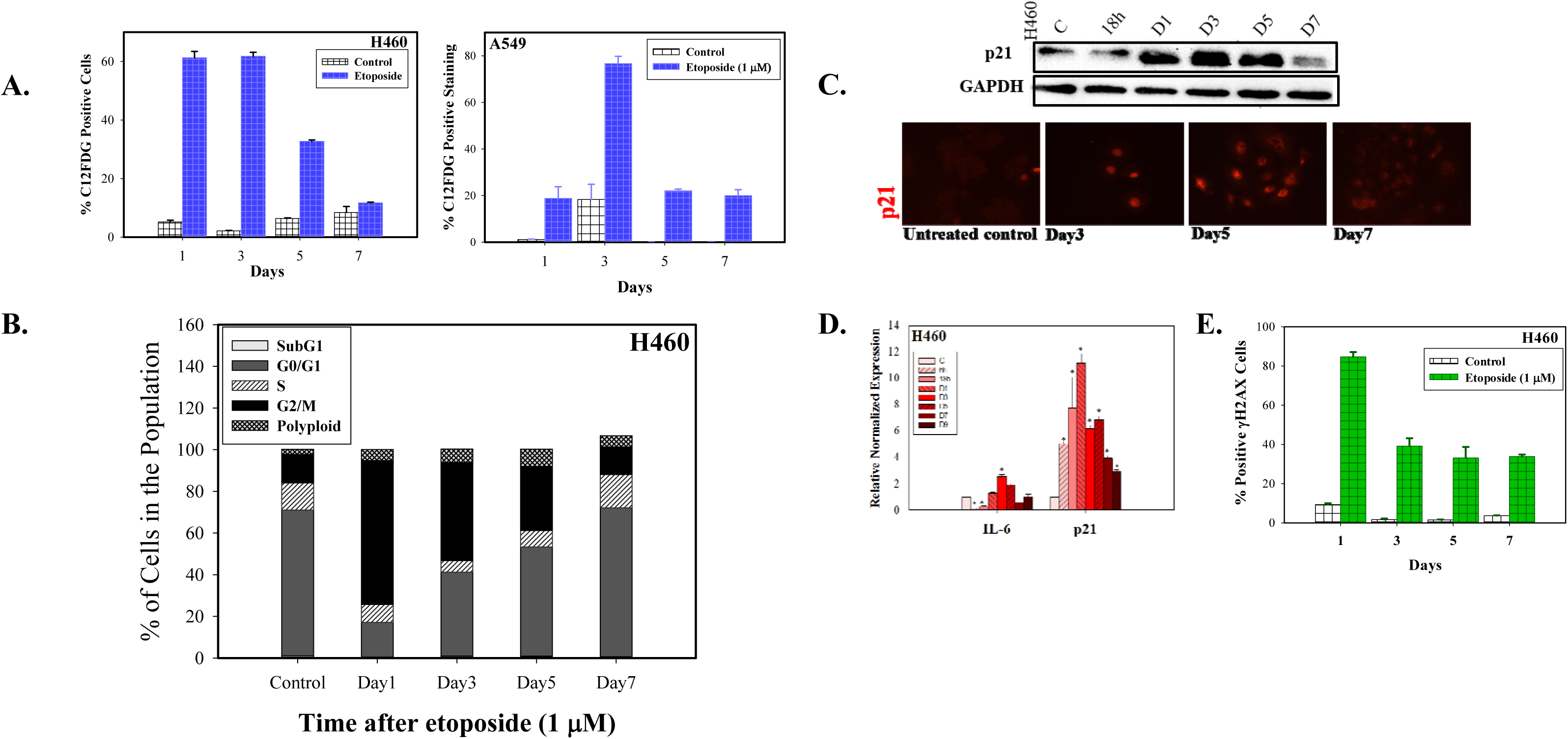
Proliferative recovery is associated with a decline in SA- β-galactosidase, entry into G1 and reduction in p21^Waf1^ expression. **A.** C_12_FDG staining of H460 and A549 cells following 24 h exposure to 1 μM etoposide followed by FACS analysis over the indicated time points (n=3). **B.** Cell cycle distribution after exposure of H460 cells to 1 μM etoposide (n=3). **C.** Western blot showing the induction and decline of p21^Waf1^ over time in H460 cells following 1μM etoposide and immunofluorescence labeling for p21^Waf1^. DNA stained with DAPI (20x objective) (n=2). **D**. qRT-PCR to monitor expression of IL-6 and *p21^Waf1^* genes following 1 μM etoposide exposure (n=2). **E.** Induction and repair of DNA damage in etoposide-treated H460 cells based on intensity of phosphorylated y-H2AX fluorescence quantified by flow cytometry. (n=3). Data are expressed as standard error of the mean (SEM) for “n” independent experiments.

### Evidence for proliferative recovery based on microscopy and live cell imaging

Proliferative recovery from senescent H460 cells was further monitored by staining with crystal violet, C_12_FDG, and CFDA. As shown in **Figure 4A**, cells reverted from an enlarged, highly-vesicular (abundant cytoplasmic granules) morphology to a more rounded, smaller phenotype; the cells further demonstrated reduced C_12_FDG-staining, consistent with reversion to a state of proliferation (**Figure 4A, middle panel**). Staining with the fluorigenic filiation tracer Vybrant CFDA also demonstrated enlarged flattened cells that gave rise to proliferating cells that retained staining by the fluorigenic tracer, consistent with the premise that these are daughter cells that emerged from the senescent cell population (**Figure 4A, lower panel**).

**Figure 4.**
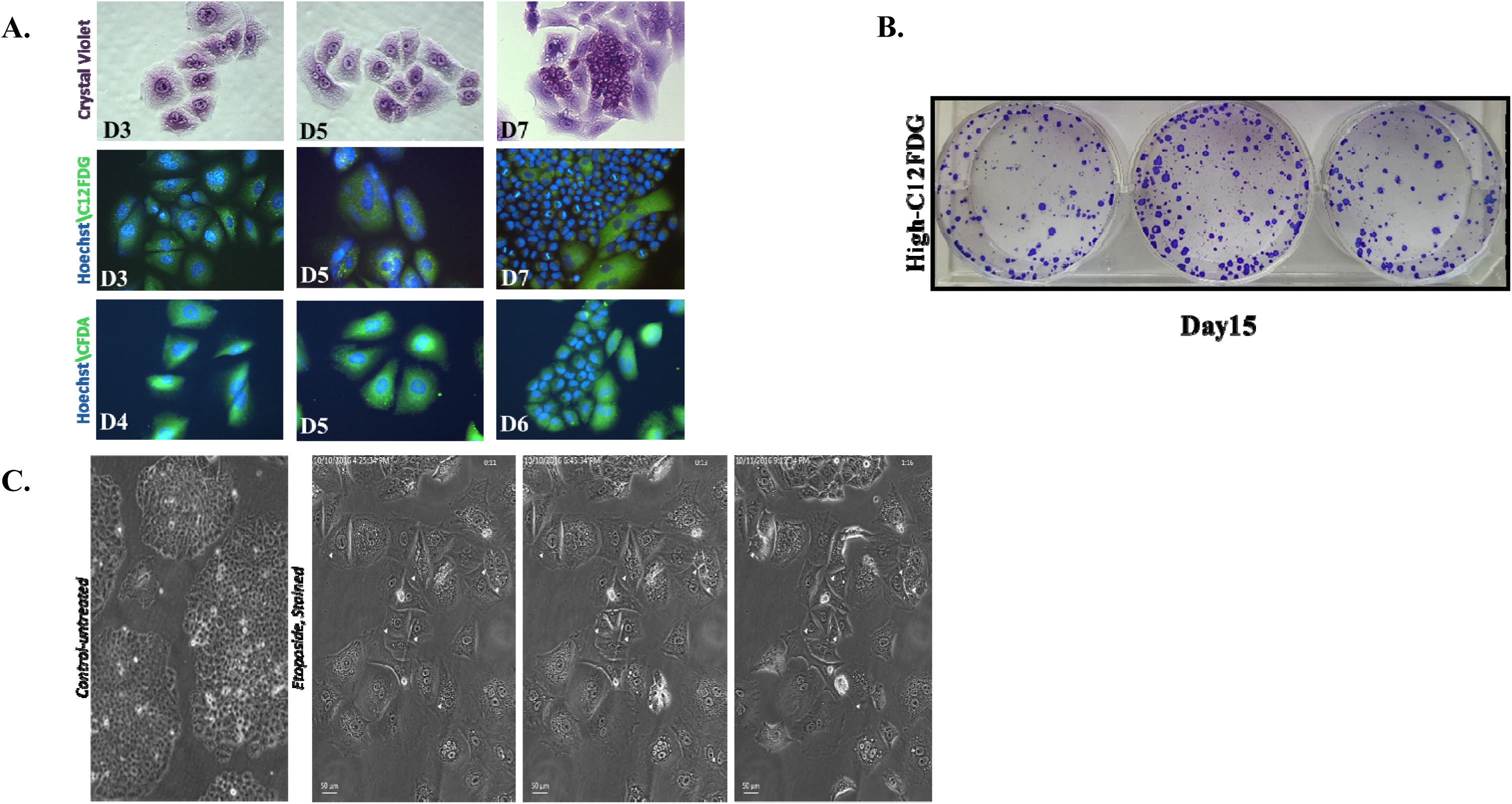
Etoposide-induced senescence in H460 cells can be reversible. **A.** H460 cells were treated with 1 μM etoposide for 24 h. *Upper panel.* Crystal violet on days 3, 5, and 7. *Middle panel.* C_12_FDG staining on days 3, 5, and 7. *Lower panel.* Vybrant CFDA staining where cells were stained on day 3 followed by microscopy on days 4, 5, and 6. DNA stained with Hoechst 33342. Fields are representative of separate experimental events. (20 x objective) (n=2) **B.** High-C_12_FDG positive cells were reseeded and monitored for colony formation. Colonies stained with crystal violet at day 15 after etoposide exposure (day 12 after sorting). **C.** Time-lapse live-cell microscopy of H460 high C12FDG-positive cells performed for 40 h, 5 days following flow cytometry-based sorting. Arrows indicate mitotic events. Data are expressed as standard error of the mean (SEM) for “n” independent experiments.

Since it is possible that the proliferating cells were not derived solely from the senescent population, cells were labeled with C_12_FDG followed by flow cytometric cell sorting and subsequent seeding of the highest C_12_FDG-stained subpopulations^15^. As shown in **Figure 4B**, the highly C_12_FDG-positive cells demonstrated restoration of proliferative capacity.

While the data generated in H460 mass cultures and senescence-enriched populations are consistent with TIS being reversible, we recognized that a careful lineage tracing approach is essential to distinguish between cancer cells that might have initially evaded etoposide-induced senescence from cells that had undergone TIS, but recovered proliferative potential. Consequently, the high C_12_FDG-positive population was reseeded and monitored by live cell imaging using phase contrast microscopy. The temporal response after etoposide-induced senescence was found to be quite heterogeneous. Over the period of observation (**Supplementary Videos 1, 2, and 3**), the majority of the senescent cell population persisted in a growth-arrested state, while a few cells were observed to eventually surrender to stress and switch to apoptosis (or possibly mitotic catastrophe). However, occasional hypertrophic, flattened senescent cells that had remained in an arrested state were found to be capable of recovering the ability to divide (**Figure 4C**). Whether this is simply a stochastic outcome or reflective of unique characteristics of subsets in the tumor cell population, is a question that we will attempt to address in future work. Nevertheless, these studies, taken together with the data in **Figure 4**, appear to establish the reversibility of chemotherapy-induced senescence.

## Discussion

Although the bulk of the literature has focused on the cytoprotective function(s) of autophagy, we and others have shown that in a number of studies interference with autophagy failed to alter drug or radiation sensitivity or to promote apoptosis^15,27,28^, as is also the case in the current work. In the case of radiation, the non-cytoprotective function of autophagy was dependent on the cells being mutant or null in p53 ^27^. However, in the current work, it is clear that autophagy induced by etoposide in the H460 cells (p53 WT) is *non-cytoprotective,* an observation we were able to replicate with other cytotoxic agents (data not shown). These findings are consistent with a recent report by Eng et al^35^ demonstrating non-cytoprotective autophagy induced by more than 30 chemotherapeutic drugs in the A549 NSCLC cell line. Etoposide-induced non-cytoprotective autophagy did not appear to affect the profile of growth arrest and proliferative recovery, indicating that this form of autophagy is unlikely to contribute to drug sensitivity.

Studies of oncogene-induced senescence have been suggestive of a close relationship between autophagy and senescence^36,37^. However, the relationship has not been consistent in different experimental models^38,39^. In both this work as well as our own previous studies^4,5,15^, it is quite clear that autophagy and senescence are dissociable.

On the other hand, senescence induction and resolution showed a more consistent pattern with growth arrest and recovery. Our laboratories previously reported on the promotion of senescence in response to chemotherapy and radiation, and furthermore provided evidence that therapy-induced senescence (TIS) may, in fact, be reversible ^14–16,40^. Other research groups have also published data supporting the premise that TIS may be reversible^17–26^. The most significant finding generated by the current work, utilizing multiple and complementary approaches, is that recovery from senescence can and does occur even from a highly β-galactosidase-active population independently of autophagy; this finding has been reproduced in breast tumor cells exposed to doxorubicin (manuscript in preparation). Another important observation is that the fate of senescent cells following a single exposure to etoposide is largely heterogeneous; as is evident from the live time-dependent imaging studies, the bulk of senescent cells remain in a growth-arrested state, a few isolated cells fail to sustain cellular stress associated with DNA damage and surrendered to cell death, while a subpopulation of senescent cells recover proliferative capacity.

A number of factors may be responsible for allowing proliferative recovery to occur in this experimental system. One is the absence of functional p16^INK4a^ in these cells^41^. While the activation of the p53-p21 axis accompanied by the senescence-associated secretory phenotype (SASP) may be sufficient to induce senescence-associated growth arrest in response to etoposide-induced DNA damage, a sustained senescence-based growth arrest is likely to be dependent upon p16^INK4a^ function^24,42^. Since inactivation or loss of p16 is quite common in cancer cells ^43,44^, reversibility of chemotherapy-induced senescence might not be an infrequent observation. The p16^INK4a^-retinoblastoma protein (RB) pathway has also been shown to be pivotal to the formation of Senescence-associated Heterochromatic Foci (SAHF), which are absent in this model (data not shown). SAHF might contribute to maintenance of the senescent phenotype^45^.

Taken together with our previous findings relating to regrowth following radiation-induced senescence, these observations, based on enrichment of senescence and real-time microscopy, support the premise that senescence induced by chemotherapy or radiation can be reversed in tumor cells and this reversal is independently of autophagy. If this proves to be the case in tumor-bearing animals, we can speculate that senescence could be the basis for some forms of tumor dormancy. That is, it is possible to imagine that after being exposed to cycles of cytotoxic chemotherapy, a few senescent cancer cells can reside in a dormant state at distant sites and eventually resume proliferation, contributing to or being responsible for disease recurrence.

Footnote1: We were unable to detect Senescence-associated Heterochromatic Foci (SAHF), relying on histone H3 trimethylation lysine 9 (H3K9me3) as a classic senescence-associated histone modification^46^ (data not shown). These findings are consistent with the apparent absence of global structural changes in chromatin in DAPI-stained nuclei. It is apparently not unusual that H3K9me3-positive nuclear foci may not be evident in models of DNA damage-induced senescence^44,45^.

## Methods

### Reagents, drugs and antibodies

Etoposide, or VP-16-213, Sigma-Aldrich (E1383), lipofectamine (Invitrogen, 11668-019), Puromycin (Sigma-Aldrich, P8833), C_12_FDG (Life Technologies, D2893), Hoechst 33342 (ThermoFisher Scientific, H1399), Vybrant CFDA (Thermo Fisher, V12883). Primary antibodies: SQSTM1/p62 (BD Biosciences, 610497), *ATG5* (Cell Signaling Technology, 2630), LC3B (Cell Signaling Technology, 3868), TP53 (BD Biosciences, 554293), Cip1/p21 (BD Biosciences, 610234), GAPDH (Cell Signaling Technology, 2118). Conjugate antibodies: γ2AX antibody (BD Biosciences, 560445).

### Cell culture and drug exposure

The wild-type (WT) TP53 H460 and A549 non-small cell lung cancer cell lines were generously provided by Dr. Richard Moran and Dr. Charles Chalfant, respectively, at Virginia Commonwealth University. The ATG5-knocked down H460 variant was generated in our laboratory: mission shRNA bacterial stocks for *ATG5* were purchased from Sigma-Aldrich (TRCN00151963) and lentivirus generation was conducted in the HEK 293TN cells. The process involved cotransfection with lipofectamine with a packaging mixture of psPAX2 and pMD2.G constructs (Addgene, 12260, 12259). After 48 h, viruses shed into the media were collected and used to infect H460 cells under ultrasonic centrifugation for 2 h. Selection was performed in Puromycin (1-2 μg/ml).

All cells were cultured in DMEM supplemented with 10% (v/v) fetal bovine serum (Thermo Scientific, SH30066.03), 100 U/ml penicillin G sodium (Invitrogen, 15140-122), and 100 μg/ml streptomycin sulfate (Invitrogen, 15140-122). Cells were maintained at 37°C under a humidified, 5% CO2 atmosphere at subconfluent densities.

At all etoposide concentration utilized, cells were exposed to the drug-containing medium for 24 h, followed by replacement with fresh medium. Incubation with Chloroquine (CQ, 10 μM) or Bafilomycin A1 (Baf, 5 nM) were the pharmacological approaches utilized to interfere with lysosomal acidification and autophagosome/lysosome fusion, respectively. Cells were treated with the autophagy inhibitors for 3 h prior to the subsequent exposure to both etoposide and the autophagy inhibitor for an additional 24 h to ensure sufficient blockade of autophagy. Drugs were protected from light during handling.

### Growth inhibition and clonogenic survival

Growth curves were generated by Trypan blue exclusion. Cells were seeded, treated (on day 0), and counted at the indicated time points following the removal of the drug from the medium or after sorting. For the clonogenic assay, cells were seeded, pre-treated with CQ (10 μM) or Baf (5 nM) for 3 h, then treated with etoposide (0.25, 0.5, 1.0, or 5 μM) alone or in combination with CQ or Baf; drugs were removed and replaced with fresh media after 24 h. Cells were incubated for 7 days, then fixed with methanol, stained with crystal violet, and counted (ColCount, Discovery Technology International).

### Flow cytometry and fluorescence microscopy

All of the flow cytometry analyses were performed using BD FACSCanto II and BD FACSDiva software at the Virginia Commonwealth University Flow Cytometry Core Facility. For C_12_FDG, acridine orange, Annexin V/Propidium Iodide, γH2AX, and cell cycle analyses, 10,000 cells per replicate within the gated region were analyzed. Three replicates for each condition were analyzed in each independent experiment. Labeling procedures, gating, and analysis followed our previously published protocols with minor adjustment for the tested cell line^5,15,47^.

**Evaluation of senescence by β-galactosidase and C_12_FDG staining**

β-galactosidase labeling was performed as previously described by Dimri et al^48^ and in our previous publications^5,15,47^. Phase contrast images were taken using an Olympus inverted microscope (20X objective, Q-Color3™ Olympus Camera; Olympus, Tokyo, Japan). The C12FDG staining protocol was adopted from Debacq-Chainiaux et al^30^. Cells were initially treated with Baf (100 nM) for 1 h to accomplish lysosomal alkalization, followed by incubation with C_12_FDG in complete media for 2 h at 37°C. Cells were harvested for flow cytometry and imaging (20X objective, Q-Color3™ Olympus Camera, 488 filter, Olympus, Tokyo, Japan).

### Western blotting and immunofluorescence

Western blots were performed as previously described^47^. Primary antibodies were used at a 1: 1000 dilution except for GAPDH (1:8000 dilution). For immunofluorescence, at each time point, cells were fixed with methanol, permeabilized with 0.1% Triton X-100, and blocked with 1% bovine serum albumin (BSA). Cells were exposed to a 1:100 dilution of p21 antibodies and incubated overnight at 4°C, followed by exposure to the secondary antibody for 1 h at room temperature. After incubation, cells were mounted with DAPI and imaging was performed with an Olympus inverted microscope (20X objective, Q-Color3™ Olympus Camera, 555 filter and UV light, Olympus, Tokyo, Japan).

### Promotion of apoptosis by Annexin V/ propidium iodide staining and TUNEL assay

Quantification of apoptotic cells via flow cytometry was achieved per the manufacturer’s instructions (AnnexinV-FITC apoptosis detection kit, BD Biosciences, 556547). For the terminal deoxynucleotidyl transferase dUTP nick end labeling (TUNEL) assay, adherent cells were harvested and centrifuged at 10,000 rpm for 5 min onto slides (Shandon Cytospin 4, Thermal Electron Corp) 48 h after etoposide removal. Slides were fixed with 4% formaldehyde for 10 min and then washed with PBS for 5 min at room temperature. The cells were then fixed with a 1:2 dilution of acetic acid and ethanol for 5 min, followed by staining with a 1:1,000 dilution of DAPI at room temperature. Coverslips were sealed using clear nail polish and apoptosis was assessed by evaluating three fields per condition with an Olympus inverted microscope (20X objective, Q-Color3™ Olympus Camera, 488 filter and UV light, Olympus, Tokyo, Japan).

### Cell Cycle Distribution by Propidium iodide (PI)

Cells were harvested with 0.1% trypsin and neutralized with medium. After centrifugation, the cells were washed with PBS, then resuspended in a PI solution [50 μg/ml PI, 4 mM sodium citrate, 0.2 mg/ml DNase-free RNase A, and 0.1% Triton-X 100] for 1 h at room temperature, while being protected from light. Before analysis, NaCl was added to the cell suspensions to achieve a final concentration of 0.20 M. The cell suspensions were then analyzed by flow cytometry.

### Vybrant CFDA Cell Tracking

Cells were stained with a 1% PBS solution containing Vybrant CFDA at a final concentration of 25 μM for 30 min. At the indicated time points, imaging was performed using an Olympus inverted microscope (20X objective, Q-Color3^TM^ Olympus Camera, 555 filter and UV light, Olympus, Tokyo, Japan).

### qRT-PCR

RNA was isolated from cell cultures using an RNAqueous Micro Total RNA Isolation kit (Thermo Fisher Scientific) according to the manufacturer’s recommendations. RT-PCR was carried out using the RETROscript kit (Ambion) to generate cDNA, followed by a standard SYRB^®^ green-based PCR assay, as described previously (Elmore et al.,^49^ and Sachs et al.,^50^). The 2-∆∆Ct method was used to determine relative expression of the two senescence-associated genes.

### Live cell imaging

Time lapse imaging of cells in culture was performed using a Zeiss Cell Observer microscope (Carl Zeiss Microscopy, Thornwood, NY), equipped with a Pecon stage incubator (set to 37**°** C, 5% CO_2_), an Axiocam MRm camera, and a Prior motorized XY stage (programmed to re-visit multiple selected sites in the culture dish). Phase contrast images were collected using a 10x / 0.3 NA Plan-Neofluar objective lens at 5 min intervals over a 48 h period. Approximately 10 fields of view were time-lapse imaged in the culture dish for each experimental trial. Images were collected and processed using Zen software.

**Supplementary Videos 1, 2 and 3.** H460 cells were exposed to etoposide (1μM for 24 hours) and then sorted by flow cytometry based on C_12_FDG staining. Only the high C12FDG-positive cells were reseeded and monitored for 40 h, 5 days following flow cytometry-based sorting. Senescent cells exhibited heterogeneous fates; where many remained arrested, some restored mitotic capacity.

## Acknowledgments

This work was supported by the Office of the Assistant Secretary of Defense for Health Affairs through the Breast Cancer Research Program [grant no. W81XWH-14-1-0088 (DAG)], Massey Center Support Grant P30 CA016059 and NIH grant # RCA206028, The authors would like to thank Dr. Khushboo Sharma for assistance with generating the shControl, shAtg5 and GFP-LC3 H460 cells and Dr. Moureq Alotaibi for establishing the C_12_FDG-based cell sorting protocol with the help of Julie Farnsworth.

